# Evidence of exposure to SARS-CoV-2 in cats and dogs from households in Italy

**DOI:** 10.1101/2020.07.21.214346

**Authors:** E.I. Patterson, G. Elia, A. Grassi, A. Giordano, C. Desario, M. Medardo, S.L. Smith, E.R. Anderson, T. Prince, G.T. Patterson, E. Lorusso, M.S. Lucente, G. Lanave, S. Lauzi, U. Bonfanti, A. Stranieri, V. Martella, F. Solari Basano, V.R. Barrs, A.D. Radford, U. Agrimi, G. L. Hughes, S. Paltrinieri, N. Decaro

**Author notes:** Correspondence to: Nicola Decaro, DVM, PhD, Dipl. ECVM, Department of Veterinary Medicine, University of Bari, Valenzano, Bari, Italy.

## Abstract

SARS-CoV-2 originated in animals and is now easily transmitted between people. Sporadic detection of natural cases in animals alongside successful experimental infections of pets, such as cats, ferrets and dogs, raises questions about the susceptibility of animals under natural conditions of pet ownership. Here we report a large-scale study to assess SARS-CoV-2 infection in 817 companion animals living in northern Italy, sampled at a time of frequent human infection. No animals tested PCR positive. However, 3.4% of dogs and 3.9% of cats had measurable SARS-CoV-2 neutralizing antibody titers, with dogs from COVID-19 positive households being significantly more likely to test positive than those from COVID-19 negative households. Understanding risk factors associated with this and their potential to infect other species requires urgent investigation.

**One Sentence Summary:** SARS-CoV-2 antibodies in pets from Italy.

## Main Text

Severe acute respiratory syndrome coronavirus 2 (SARS-CoV-2) emerged in late December 2019 in Wuhan, Hubei province, China (1), possibly as a spillover from bats to humans (2), and rapidly spread worldwide becoming a pandemic (3). Although the virus is believed to spread almost exclusively by human-to-human transmission, there are concerns that some animal species may contribute to the ongoing SARS-CoV-2 pandemic epidemiology (4). To date, sporadic cases of SARS-CoV-2 infection have been reported in dogs and cats. These include detection of SARS-CoV-2 RNA in respiratory and/or fecal specimens of dogs and cats with or without clinical signs (5-7), as well as of specific antibodies in sera from pets from coronavirus disease 2019 (COVID-19) affected areas (7,8). In addition, experimental infection of various animal species has demonstrated that while dogs appear poorly susceptible to SARS-CoV-2 infection, developing asymptomatic infections and shedding low-titer or no virus, cats develop respiratory pathology and shed high titers of SARS-CoV-2, even being able to infect in-contact animals (9,10). Wide scale testing of susceptible species is needed to assess the extent of animal infection under more natural conditions of husbandry. Here, we conducted an extensive epidemiological survey from March to May 2020 in cats and dogs living in Italy, either in SARS-CoV-2 positive households or living in geographic areas that were severely affected by COVID-19. To our knowledge, this is the largest study to investigate SARS-CoV-2 in companion animals to date.

All animals were sampled by their private veterinary surgeon during routine healthcare visits. Sampling of animals for this study was approved by the Ethics Committee of the Department of Veterinary Medicine, University of Bari, Italy (approval number 15/2020). A total of 540 dogs and 277 cats were sampled from different Italian regions, mostly Lombardy (476 dogs, 187 cats). Animals were sampled either from regions severely affected by COVID-19 outbreaks in humans or from those that offered convenient access to samples. Oropharyngeal (306 dogs, 175 cats), nasal (185 dogs, 77 cats), and/or rectal (66 dogs, 30 cats) swabs were collected from the sampled pets. For 340 dogs and 188 cats, full signalment and clinical history were available, including breed, sex, age, exposure to COVID-19 infected humans (COVID-19 positive household, suspected COVID-19 positive household but not confirmed by specific assay, and COVID-19 negative household), presence of respiratory signs (cough, sneezing, conjunctivitis, nasal and/or ocular discharge).

Sera were available for 188 dogs and 63 cats for which complete signalment, history and location were available (Fig. 1). Additional sera were collected from diagnostic laboratories for 200 dogs and 89 cats from the affected areas, but which lacked further historical information.

**Fig. 1.**
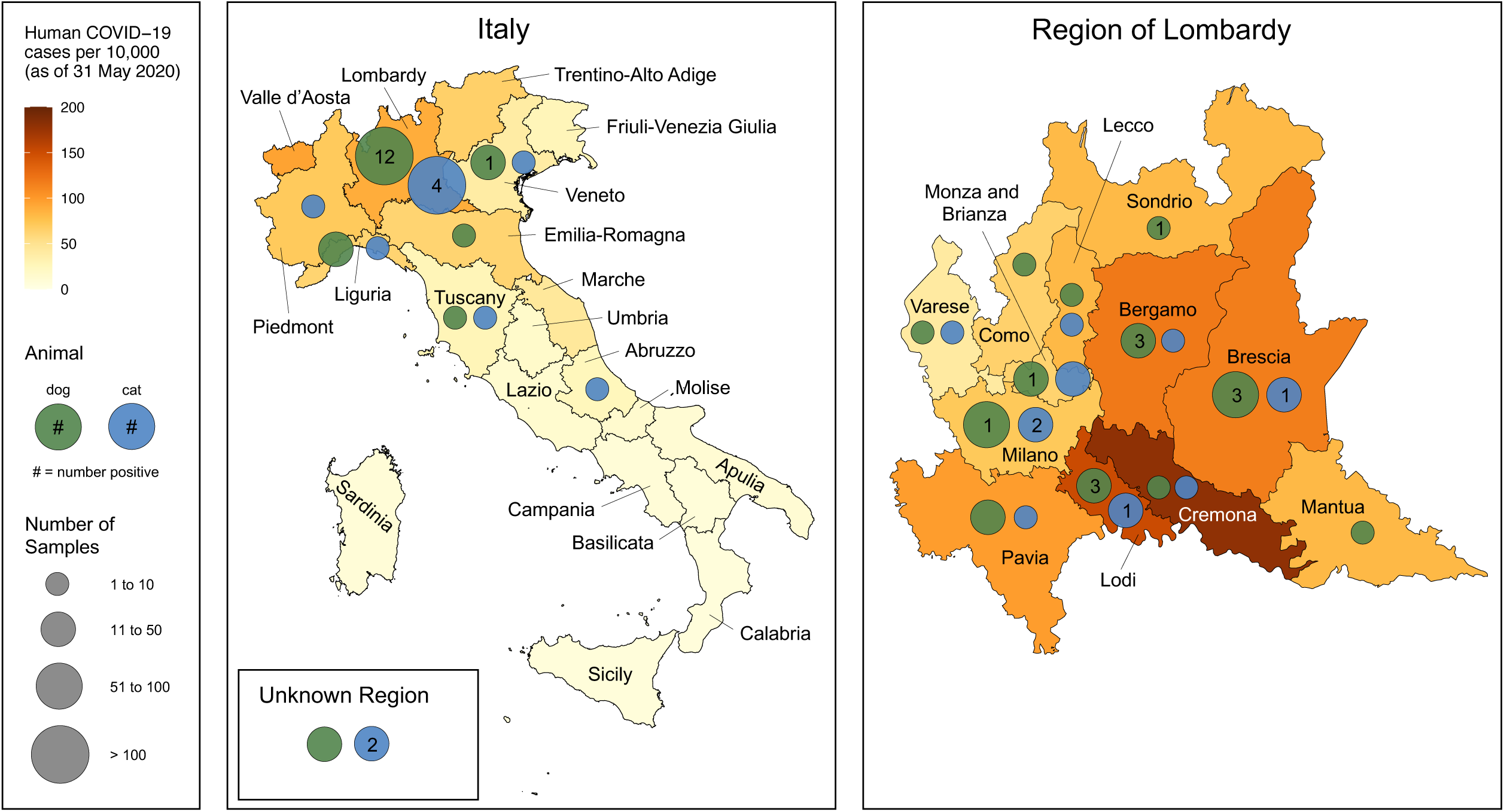
Distribution of dog and cat samples assayed for neutralizing antibody titer across Italy and the region of Lombardy. Data on human COVID-19 cases from the Italian Department of Civil Protection as of May 31, 2020 and population data from the Italian National Institute of Statistics (ISTAT), January 2019.

Detection of SARS-CoV-2 RNA used two real-time RT-PCR assays targeting nucleoprotein and envelope protein genes as previously described (11). Plaque reduction neutralization tests (PRNT) were using a previously established protocol (8) with SARS-CoV-2/human/Liverpool/REMRQ0001/2020 isolate was cultured as previously described (9). PRNT80 was determined by the highest dilution with □80% reduction in plaques compared to the control.

All of 839 collected swab samples tested negative for SARS-CoV-2 RNA, including 38 cats and 38 dogs that showed respiratory symptoms at the time of sampling, suggesting absence of active SARS-CoV-2 infection in the tested animals. In addition, 64 of these dogs and 57 of the cats that tested negative were living in households previously confirmed as having had COVID-19.

SARS-CoV-2 neutralizing antibodies were detected in 13 dogs (3.35%) and 6 cats (3.95%), with titers ranging from 1:20 to 1:160 and from 1:40 to 1:1280 in dogs and cats, respectively. Of samples from households with known COVID-19 status, neutralizing antibodies were detected in 6 of 47 dogs (12.8%) and 1 of 22 cats (4.5%) from COVID-19 positive households, 1 of 7 dogs (14.3%) and 0 of 3 cats (0%) from suspected COVID-19 positive households and 2 of 133 dogs (1.5%) and 1 of 38 cats (2.6%) from COVID-19 negative households (Table 1). For those 423 animals where an age was recorded, 0 of 30 aged less than 1 year (0%), 6 of 92 aged 1-3 years (6.5%), 3 of 102 aged 4-7 years (2.9%) and 6 of 199 aged 8 and over (3.0%) tested positive. None of the animals with neutralizing antibodies displayed respiratory symptoms at the time of sampling.

**Table 1.**
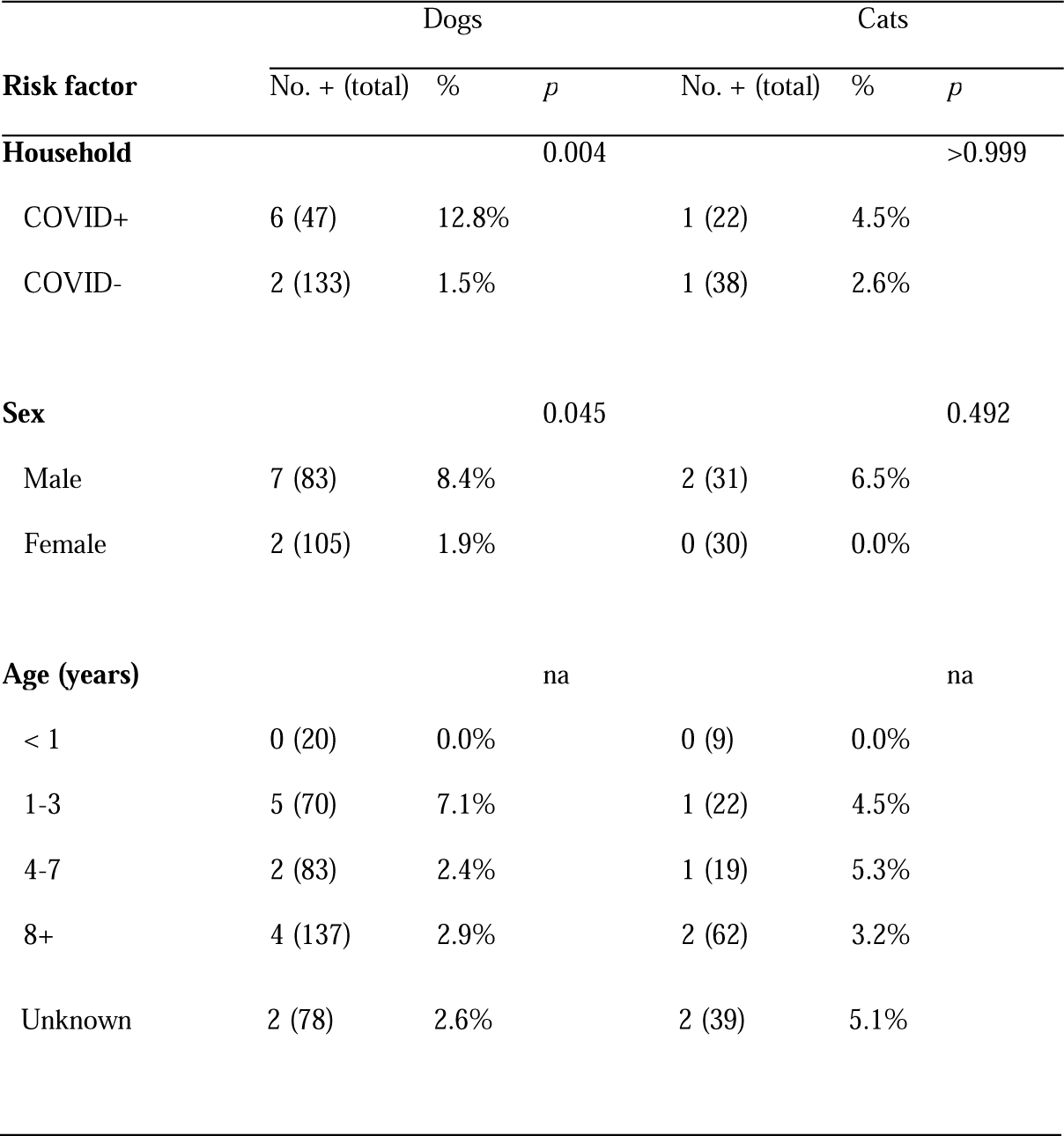
Seropositivity among dogs and cats, split into risk factor groupings where data was available. For household and sex, p value determined by Fisher’s exact test. Household COVID + defined as one or more members of a household with a confirmed positive COVID-19 test. All the information was not available for all the animals.

Reference sera or ascitic fluids from animals previously shown to be positive for canine enteric coronavirus (14), canine respiratory coronavirus (15) and feline coronavirus (16) tested negative by the PRNT assay for SARS-CoV-2, confirming the specificity of the obtained results (8).

Dogs were significantly more likely to test positive for SARS-CoV-2 neutralizing antibodies if they came from a known COVID-19 positive household (Fisher’s exact test, p=0.004) or were male (Fisher’s exact test, p=0.045). In provinces where at least 10 samples were available, there was a strong positive trend between the proportion of dogs that tested positive and the recorded burden of human disease (Spearman’s r = 0.732, p = 0.051) (Fig. 2). A similar association was observed for cats but should be viewed with caution as only four provinces met the criteria for analysis.

**Fig. 2.**
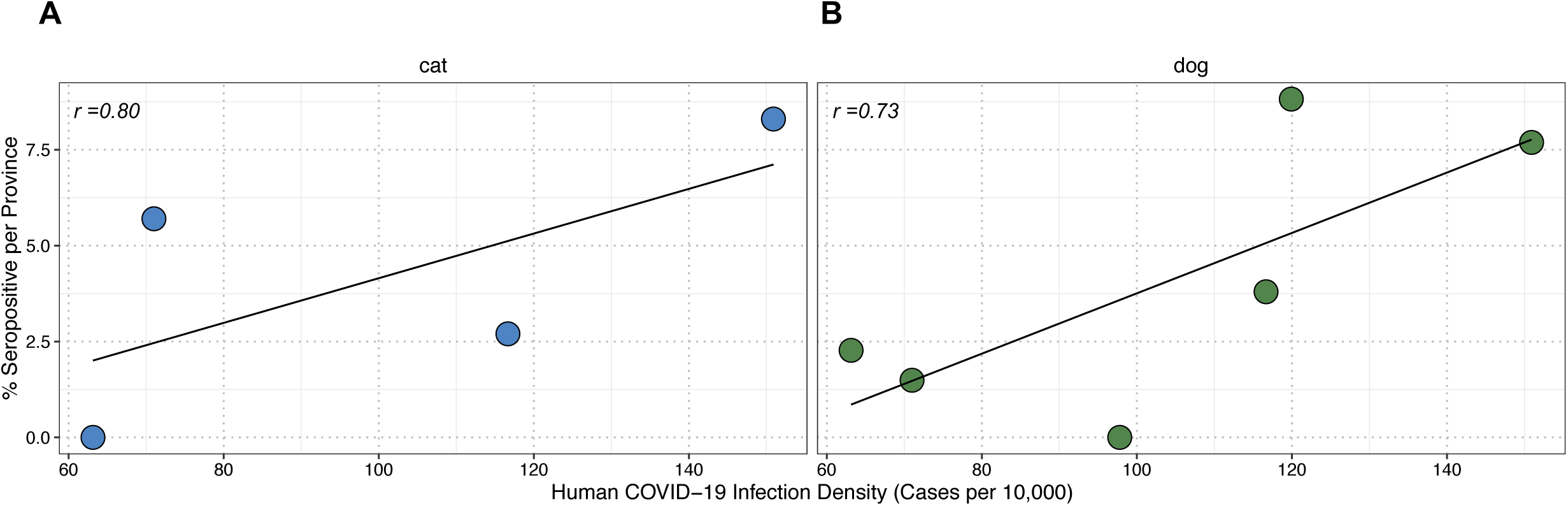
Correlation of percentage of seropositive animals per province and human COVID-19 infection density. Data points were taken from provinces with at least 10 samples. Spearman’s correlation was used to assess association.

Following its original probable transmission to humans from animals, SARS-CoV-2 has spread globally within the human population with devastating health and economic impacts. To date, SARS-CoV-2 has been sporadically detected in naturally infected dogs and cats, most of which were living in close contact with infected humans. Most studies of companion animals are small in nature, likely because of an inevitable research focus on human disease. Our results from this extensive study of SARS-CoV-2 infection in owned cats and dogs living in areas where viral transmission was active in the human population demonstrate that both cats and dogs can seroconvert under the normal conditions of pet ownership, and where the burdens of disease are highest in humans.

The link between SARS-CoV-2 household infection and a pet’s seropositivity was only apparent for dogs, possibly suggesting greater interaction between positive people and their household dogs as compared to cats. This contrasts experimental studies where dogs were less susceptible to infection (9). In addition, a higher proportion of male dogs were seropositive compared to female dogs. Future studies in animals and humans should investigate whether this phenomenon is based in physiological or behavioral differences between males and females. Although there are clear gender differences in outcomes in human COVID-19 infections, with males at higher risk of severe disease, there seems to be no evidence for a difference in infection risk (17). None of the 30 juvenile animals, less than one year-of-age, tested positive. Our findings are consistent with reports of other seropositive naturally exposed cats and dogs which were all adult (6, 7). These findings support use of older animals in experimental infections, which are currently performed on animals less than one year-of-age (9) and may therefore underestimate SARS-CoV-2 susceptibility.

In contrast to the serology results, all animals tested negative by PCR, including those animals living in households with confirmed COVID-19 human infection and those with and without respiratory symptoms. This suggests that whilst pet animals can seroconvert, they may shed virus for relatively short periods of time. In experimental studies, cats stopped shedding virus by 10 days post infection (dpi) and developed neutralizing antibody responses by 13 dpi (9). Similar results were reported in experimental infection of dogs, in which virus was detected in faeces up to 6 dpi, but not in oropharyngeal swabs (6). However, in a naturally infected Pomeranian dog SARS-CoV-2 RNA was detected from nasal swabs by quantitative RT-PCR for at least 13 days at low titer, whilst the virus was not detected in faecal/rectal samples (7), suggesting that virus shedding patterns may vary in some animals. Half of the challenged dogs had detectable antibodies by 14 dpi. These studies and our own highlight similar challenges in detecting SARS-CoV-2 infection that exist for both humans and animals (18). It is not possible with our field data set to estimate the time of infection in animals that were seropositive, and restrictions on human and animal movement during the pandemic may have delayed visits to veterinary practitioners where sampling occurred. We advocate the inclusion of pets in ongoing assessments of community and household shedding to improve detection of active infection.

In this extensive epidemiological survey of SARS-CoV-2, we found that companion animals living in areas of high human infection can become infected. Our results suggest that dogs warrant further investigation regarding SARS-CoV-2 susceptibility in contrast to experimental studies which suggested cats were most susceptible (9). We also observed seropositivity rates in animals comparable to those of humans via community sampling at a similar time in European countries (19-21). This suggests that infection in companion animals is not unusual. Based on current knowledge, it is unlikely that infected pets play an active role in SARS-CoV-2 transmission to humans. However, animal-to-human transmission may be more likely under certain environmental conditions, such as the high animal population densities encountered on infected mink farms (22). As and when human transmission becomes rarer and contact tracing becomes more accessible, serological surveillance of pets may be advocated to develop a wholistic picture of community disease dynamics and ensure that all transmission opportunities are terminated.

## Acknowledgments

SARS-CoV-2 RNA extracts from infected human patients that were used as positive controls in the real-time RT-PCR assays were kindly provided by Istituto Zooprofilattico della Puglia e della Basilicata, Foggia, and by Istituto Zooprofilattico dell’Abruzzo e del Molise “G. Caporale”, Teramo, Italy. We are grateful to the vets that contributed to the sample collection in the COVID-19 severely affected Italian regions.

## Funding

This work was supported by grants of Fondazione CARIPLO - Misura a sostegno dello sviluppo di collaborazioni per l’identificazione di terapie e sistemi di diagnostica, protezione e analisi per contrastare l’emergenza Coronavirus e altre emergenze virali del future, project “Genetic characterization of SARS-CoV2 and serological investigation in humans and pets to define cats and dogs role in the COVID-19 pandemic (COVIDinPET)”. EIP was supported by the Liverpool School of Tropical Medicine Director’s Catalyst Fund award. SLB and ADR were supported by the DogsTrust. GLH was supported by the BBSRC (BB/T001240/1), the Royal Society Wolfson Fellowship (RSWF\R1\180013), NIH grants (R21AI138074 and R21AI129507), UKRI (20197) and the NIHR (NIHR2000907). GLH and TP are affiliated to the National Institute for Health Research Health Protection Research Unit (NIHR HPRU) in Emerging and Zoonotic Infections at University of Liverpool in partnership with Public Health England (PHE), in collaboration with Liverpool School of Tropical Medicine, the University of Oxford, and the University of Liverpool. GLH is based at LSTM. The views expressed are those of the author(s) and not necessarily those of the NHS, the NIHR, the Department of Health or Public Health England.

## Author contributions

EIP and ND designed the study and wrote the first draft of the manuscript; SLS, ADR, TP, GTP and GLH edited the manuscript, GE, AG and SP contributed to the study’s design, CD, SLS, ERA, TP, EL, MSL, and GL performed experiments; AG, MM, AS, SL, UB, VM, FSB, VRB, ADR, UA, GTP and GLH analyzed data.

## Competing interests

Authors declare no competing interests; and

## Data and materials availability

All data is available in the main text or the supplementary materials.

## Materials and Methods

### Samples

All animals were sampled by their private veterinary surgeon during a healthcare visit for other reasons. A total of 540 dogs and 277 cats were sampled from different Italian regions, mostly Lombardy (476 dogs, 187 cats). Animals were sampled either from regions severely affected by COVID-19 outbreaks in humans or from those that offered convenient access to samples. Oropharyngeal (306 dogs, 175 cats), nasal (185 dogs, 77 cats), and/or rectal (66 dogs, 30 cats) swabs were collected from the sampled pets. For 340 dogs and 188 cats, full signalment and clinical history were available, including breed, sex, age, exposure to COVID-19 infected humans (COVID-19 positive household, suspected COVID-19 positive household but not confirmed by specific assay, and COVID-19 negative household), presence of respiratory signs (cough, sneezing, conjunctivitis, nasal and/or ocular discharge).

Sera were available for 188 dogs and 63 cats for which complete signalment, history and location were available (Figure 1). Additional sera were collected from diagnostic laboratories for 200 dogs and 89 cats from the affected areas, but which lacked further historical information.

Sampling of animals for this study was approved by the Ethics Committee of the Department of Veterinary Medicine, University of Bari, Italy (approval number 15/2020).

### Polymerase chain reaction

Sample preparation and RNA extraction were carried out in the biosafety level 3 containment laboratory at the Department of Veterinary Medicine, University of Bari, Italy. Detection of SARS-CoV-2 RNA used two real-time RT-PCR assays targeting nucleoprotein and envelope protein genes as previously described *(11)*.

### Plaque reduction neutralization test (PRNT)

The SARS-CoV-2/human/Liverpool/REMRQ0001/2020 isolate was cultured in Vero E6 cells as previously described (12). PRNTs were performed as previously described (13). Briefly, sera were heat inactivated at 56°C for 1 hour and stored at -20°C until use. Dulbecco’s minimal essential medium (DMEM) containing 2% fetal bovine serum (FBS) and 0.05 mg/mL gentamicin was used for serial two-fold dilutions of serum. SARS-CoV-2 at 800 PFU/mL was added to an equal volume of diluted serum and incubated at 37°C for 1 hour. The virus-serum dilution was inoculated onto Vero E6 cells, incubated at 37°C for 1 hour, and overlaid as in standard plaque assays. Cells were incubated for 48 hours at 37°C and 5% CO_2_ then fixed with 10% formalin and stained with 0.05% crystal violet solution. PRNT_80_ was determined by the highest dilution with 80% reduction in plaques compared to the control. Samples with detectable neutralizing antibody titer were repeated as technical replicates for confirmation.

### Data analysis

Fisher’s exact test was used to analyze differences in antibody detection from households with known COVID-19 infection status, and antibody detection from male and female animals. Spearman correlation was used to analyze the relationship between human COVID-19 case numbers and detection of antibodies in animals. All statistical analyses were performed in GraphPad Prism.

## Notes

### Competing Interest Statement

The authors have declared no competing interest.

## References and Notes

1. P. Zhou, X. L. Yang, X. G. Wang, B. Hu, L. Zhang, W. Zhang, H. R. Si, Y. Zhu, B. Li, C. L. Huang, H. D. Chen, J. Chen, Y. Luo, H. Guo, R. D. Jiang, M. Q. Liu, Y. Chen, X. R. Shen, X. Wang, X. S. Zheng, K. Zhao, Q. J. Chen, F. Deng, L. L. Liu, B. Yan, F. X. Zhan, Y. Y. Wang, G. F. Xiao, Z. L. Shi, A pneumonia outbreak associated with a new coronavirus of probable bat origin. Nature 579, 270–273 (2020).

2. N. Decaro, A. Lorusso, Novel human coronavirus (SARS-CoV-2): a lesson from animal coronaviruses. Vet. Microbiol. doi: 10.1016/j.vetmic.2020.108693 (2020).

3. World Health Organization, https://www.who.int/emergencies/diseases/novel-coronavirus-2019/situation-reports

4. T. McNamara, J. A. Richt, L. Glickman, A critical needs assessment for research in companion animals and livestock following the pandemic of COVID-19 in humans. Vector Borne Zoonotic Dis. doi: 10.1089/vbz.2020.2650 (2020).

5. A.S. Abdel-Moneim, E. M. Abdelwhab. Evidence for SARS-CoV-2 Infection of Animal Hosts. Pathogens. 30;9(7):E529. doi: 10.3390/pathogens9070529 (2020).

6. A. Newman, D. Smith, R. R. Ghai, R. M. Wallace, M. K. Torchetti, C. Loiacono, L. S. Murrell, A. Carpenter, S. Moroff, J. A. Rooney, C. B. Behravesh. First Reported Cases of SARS-CoV-2 Infection in Companion Animals - New York, March-April 2020. Morb Mortal Wkly Rep. 69(23):710–713 (2020).

7. T. H. C. Sit, C.J. Brackman, S. M. Ip, K.W.S. Tam, P.Y.T. Law, E.M.W. To, V.Y.T. Yu, L.D. Sims, D.N.C. Tsang, D.K.W. Chu, R.A.P.M Perera, L.L.M. Poon, M. Peiris. Infection of dogs with SARS-CoV-2. Nature. doi: 10.1038/s41586-020-2334-5 (2020).

8. Q. Zhang, H. Zhang, K. Huang, Y. Yang, X. Hui, J. Gao, X. He, C., Li, W. Gong, Y. Zhang, C. Peng, X. Gao, H. Chen, Z. Zou, Z Shi, M. Jin, SARS-CoV-2 neutralizing serum antibodies in cats: a serological investigation. bioRxiv doi: 10.1101/2020.04.01.021196 (2020).

9. J. Shi, Z. Wen, G. Zhong, H., Yang, C. Wang, R. Liu, X. He, L. Shuai, Z. Sun, Y. Zhao, L. Liang, P. Cui, J. Wang, X. Zhang, Y. Guan, H. Chen, Z. Bu, Susceptibility of ferrets, cats, dogs, and different domestic animals to SARS-coronavirus-2. Science doi: 10.1126/science.abb7015 (2020).

10. P. J. Halfmann, M. Hatta, S. Chiba, T. Maemura, S. Fan, M. Takeda, N. Kinoshita, S. Hattori, Y. Sakai-Tagawa, K. Iwatsuki-Horimoto, M. Imai, Y. Kawaoka. Transmission of SARS-CoV-2 in Domestic Cats. N Engl J Med. doi: 10.1056/NEJMc2013400 (2020).

11. V. M. Corman, O. Landt, M. Kaiser, R. Molenkamp, A. Meijer, D. K. Chu, T. Bleicker, S. Brünink S, J. Schneider, M. L. Schmidt, D. G. Mulders, B. L. Haagmans, B. van der Veer, S. van den Brink, L. Wijsman, G. Goderski, J. L. Romette, J. Ellis, M. Zambon, M. Peiris, H. Goossens, C. Reusken, M. P. Koopmans, C. Drosten, Detection of 2019 novel coronavirus (2019-nCoV) by real-time RT-PCR. Euro Surveill. 25(3). doi: 10.2807/1560-7917.ES.2020.25.3.2000045 (2020).

12. S. L. Rossi, K. E. Russell-Lodrigue, S. Z. Killeen, E. Wang, G. Leal, N. A. Bergren, H. Vinet-Oliphant, S. C. Weaver, C. J. Roy. IRES Containing VEEV vaccine protects cynomolgus macaques from IE Venezuelan equine encephalitis virus aerosol challenge. PLoS Negl. Trop. Dis. 9(5), e0003797 (2015).

13. E. I. Patterson, T. Prince, E. R. Anderson, A. Casas-Sanchez, S. L. Smith, C. Cansado-Utrilla, L. Turtle, G. L. Hughes. Methods of inactivation of SARS-CoV-2 for downstream biological assays. bioRxiv 2020.05.21.108035; doi: https://doi.org/10.1101/2020.05.21.108035 (2020).

14. N. Decaro, G. Elia, V. Martella, M. Campolo, V. Mari, C. Desario, M. S. Lucente, E. Lorusso, T. Kanellos, R. H. Gibbons, C. Buonavoglia. Immunity after natural exposure to enteric canine coronavirus does not provide complete protection against infection with the new pantropic CB/05 strain. Vaccine, 2010, 28(3), 724–729 (2010).

15. N. Decaro, C. Desario, G. Elia, V. Mari, M. S. Lucente, P. Cordioli, M. L. Colaianni, V. Martella, C. Buonavoglia. Serological and molecular evidence that canine respiratory coronavirus is circulating in Italy. Vet. Microbiol 2007, 121(3-4), 225-230 (2007).

16. E. Lorusso, V. Mari, M. Losurdo, G. Lanave, A. Trotta, G. Dowgier, M. L. Colaianni, A. Zatelli, G. Elia, D. Buonavoglia, N. Decaro. Discrepancies between feline coronavirus antibody and nucleic acid detection in effusions of cats with suspected feline infectious peritonitis. Res. Vet. Sci. 125, 421–424 (2019).

17. E. Petersen, M. Koopmans, U. Go, D. H. Hamer, N. Petrosillo, F. Castelli, M. Storgaard, S. Al Khalili, L. Simonsen. Comparing SARS-CoV-2 with SARS-CoV and influenza pandemics. Lancet Infect Dis. S1473-3099(20)30484-9 (2020).

18. S. Temmam, A. Barbarino, D. Maso, S. Behillil, V. Enouf, C. Huon, A. Jaraud, L. Chevallier, M. Backovic, P. Pérot, P. Verwaerde, L. Tiret, S. van der Werf, M. Eloit, Absence of SARS-CoV-2 infection in cats and dogs in close contact with a cluster of COVID-19 patients in a veterinary campus. bioRxiv 2020.04.07.029090 doi: 10.1101/2020.04.07.029090 (2020).

19. L. Valenti, A. Bergna, S. Pelusi, F. Facciotti, A. Lai, M. Tarkowski, A. Berzuini, F. Caprioli, L. Santoro, G. Baselli, C. Della Ventura, E. Erba, S. Bosari, M. Galli, G. Zehender, D. Prati. SARS-CoV-2 seroprevalence trends in healthy blood donors during the COVID-19 Milan outbreak. medRxiv 2020.05.11.20098442; doi: https://doi.org/10.1101/2020.05.11.20098442 (2020)

20. M. Pollan, B. Perez-Gomez, R. Pastor-Barriuso, J. Oteo, M.A. Hernan, M. Perez-Olmeda, et al. Prevalence of SARS-CoV-2 in Spain (ENE-COVID): a nationwide, population-based seroepidemiological study. The Lancet. https://doi.org/10.1016/S0140-6736(20)31483-5 (2020).

21. S. Stringhini, A. Wisniak, G. Piumatti, A.S. Azman, S.A. Lauer, H. Baysson, et al. Seroprevalence of anti-SARS-CoV-2 IgG antibodies in Geneva, Switzerland (SEROCoV-POP): a population-based study. The Lancet. https://doi.org/10.1016/S0140-6736(20)31304-0 (2020).

22. N. Oreshkova, R.J. Molenaar, S. Vreman, F. Harders, B.B. Oude Munnink, R. W. Hakze-van der Honing, N. Gerhards, P. Tolsma, R. Bouwstra, R.S. Sikkema, M.G. Tacken, M.M. de Rooij, E. Weesendorp, M.Y.. Engelsma, C.J. Bruschke, L.A. Smit, M. Koopmans, W.H. van der Poel, A. Stegeman. SARS-CoV-2 infection in farmed minks, the Netherlands, April and May 2020. Euro Surveill. 25(23). doi: 10.2807/1560-7917.ES.2020.25.23.2001005 (2020).

